# CRISPRδ: dCas13-mediated translational repression for accurate gene silencing in mammalian cells

**DOI:** 10.1101/2023.05.14.540671

**Authors:** Antonios Apostolopoulos, Hitomi Tsuiji, Yuichi Shichino, Shintaro Iwasaki

## Abstract

Current gene silencing tools based on RNA interference (RNAi) or, more recently, clustered regularly interspaced short palindromic repeats (CRISPR)⃩Cas13 systems, have critical drawbacks, such as off-target effects (RNAi) or collateral mRNA cleavage (CRISPR⃩Cas13). Thus, a more specific method of gene knockdown is needed. Here, we developed “CRISPRδ”, an approach for translational silencing, harnessing catalytically inactive Cas13 proteins (dCas13). Owing to its tight association with mRNA, dCas13 serves as a physical roadblock for scanning ribosomes during translation initiation and does not affect mRNA stability. Guide RNAs covering the start codon lead to the highest efficacy regardless of the translation initiation mechanism: cap-dependent or internal ribosome entry site (IRES)-dependent translation. Strikingly, genome-wide ribosome profiling revealed the extremely high gene knockdown specificity of CRISPRδ. Moreover, fusion of a translational repressor to dCas13 ensured further improvement of the knockdown efficacy. Our method provides a framework for translational repression-based gene silencing in eukaryotes.

## Introduction

Since the discovery of RNA interference (RNAi) ^1^, posttranscriptional gene knockdown has been a common strategy for biological research and has shown therapeutic promise ^2^. Although small RNAs used for RNAi possess 21-22 nucleotide (nt)-long sequences, their target specificity relies mainly on the complementarity of 2-8-nt-long sequences at the 5′ end ^3^. Thus, RNAi has the potential to silence “off-target” genes ^4–6^. Moreover, due to the use of a double-stranded RNA duplex as a precursor of small interfering RNA (siRNA), the induced interferon response may be another issue ^7–10^. Therefore, a method with higher accuracy and specificity has long been desired.

Recently, repurposing of a clustered regularly interspaced short palindromic repeats (CRISPR)-Cas system, a natural bacterial defense system against infecting viruses and plasmids, has allowed more specific RNA targeting. Cas13, a single type VI protein effector (class 2), associates with a guide RNA (gRNA) containing a 20-30-nt-long spacer region with a direct repeat (DR) hairpin, which anchors Cas13 proteins ^11–16^. Hybridization of the spacer region to the target RNA activates Cas13 as an RNase ^16–19^. The near-perfect complementarity between the spacer region and the target mRNA required for Cas13 activation is the basis of the superior specificity of this system compared to RNAi ^17–22^. Although the primary substrate of Cas13 should be the target RNA that is directly bound, Cas13 also degrades neighboring RNA species indiscriminately due to the solvent exposure of two catalytic higher eukaryotes prokaryotes nucleotide-binding (HEPN) domains ^11–16^. This RNase activity toward bystander RNA species (collateral activity) provides the immune function with Cas13 ^23, 24^. Earlier reports suggested that this collateral activity is limited in eukaryotic cells ^17–19^, and indeed, subsequent application of this system for targeted knockdown aligned with the goals ^20, 22, 25–42^. However, later studies have raised concerns that bystander RNA cleavage was unleashed in a wide variety of cells and led to cytotoxicity ^21, 43–53^, therefore posing an obstacle to the usage of Cas13 for transcript-specific knockdown.

However, after conversion of the catalytic residues to generate an inactive protein, Cas13 still serves as a platform to target RNA with high specificity. The tight and specific association of catalytically inert Cas13 (dead Cas13 or dCas13) can be harnessed to impede RNA-binding protein association to control alternative splicing ^19, 54^ and to track RNA mobility by GFP fusion ^17, 33, 55–57^. When fused to various effector proteins, dCas13 has also been used to induce A-to-I and C-to-U editing ^18, 48, 58–64^, m^6^A installation/removal ^65–67^, and proximity protein labeling on the defined mRNA ^68, 69^. This expanding toolkit paves the way for the spatiotemporal manipulation of target RNAs.

Here, we employed dCas13 to repress translation initiation in mammalian cells for accurate gene silencing. With the use of the RNase-inactive mutant, we reasoned that our system could circumvent the collateral activity of Cas13 in the transcriptome. Indeed, genome-wide ribosome profiling revealed the high specificity of our system for suppressing protein synthesis from the target mRNA. Tiling of the gRNAs along the target mRNA showed a steric hindrance effect of dCas13 on the preinitiation complex during the scanning process and/or a stable association at the AUG start codon, ensuring the optimal design of the gRNAs (∼23-nt length and start codon coverage). Our system applied not only to standard cap-dependent translation but also to internal ribosome entry site (IRES)-mediated translation. Moreover, fusion of the translational suppressor, eukaryotic translation initiation factor (eIF) 4E homologous protein (4EHP), further enhanced the silencing efficacy. Our method, termed CRISPRδ [delta (δ): dCas13-induced DEpLetion of Translation by blockAde], provides a useful framework for a tool to implement precise gene knockdown.

## Results

### dCas13 represses translation initiation of target transcripts

We hypothesized that the tight binding of Cas13 to an mRNA may abrogate the movement of ribosomes. Given the diversity of Cas13 family proteins ^70, 71^, we aimed to survey their potential for translational repression (Figure 1A). Considering their extensive applications for RNA biology to date ^18, 19, 48, 55–57, 59–62, 64, 65, 67–69^, we generated catalytically inactive (dCas13) proteins corresponding to four selected Cas13 proteins: *Leptotrichia wadei* Cas13a (LwaCas13a and dLwaCas13a), *Prevotella sp. P5-125* Cas13b (PspCas13b and dPspCas13b), *Porphyromonas gulae* Cas13b (PguCas13b and dPguCas13b), and *Ruminococcus flavefaciens* XPD3002 Cas13d (RfxCas13d and dRfxCas13d). As subcellular localization has been reported to affect Cas13 performance ^17, 18, 53, 65, 72^, we expressed each of those Cas13 variants with a nuclear export signal (NES) or nuclear localization signal (NLS).

**Figure 1.**
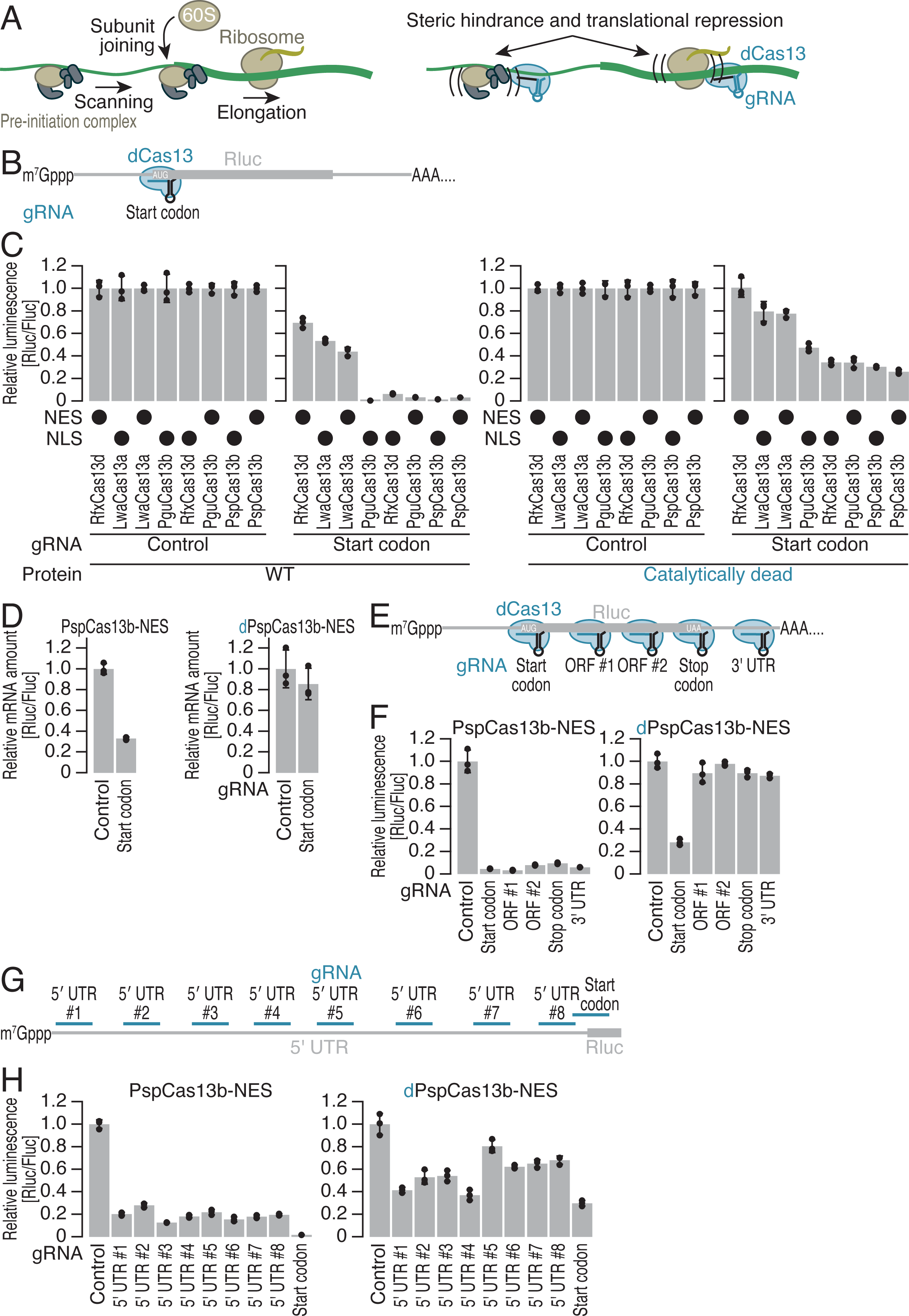
Repurposing catalytically inactive Cas13 proteins for translational repression. (A) Schematic of dCas13-induced repression of mRNA translation. dCas13 binding to an mRNA can selectively repress its translation by sterically hindering the translational machinery. (B) Schematic of the luciferase reporter and gRNA design. gRNAs targeting the start codon of Rluc reporter mRNA were designed for each of the dCas13 and Cas13 variants. A gRNA containing a nontargeting spacer sequence was used as a control. Fluc expression was used as an internal control. The start codon-targeting gRNA for PspCas13b/dPspCas13b was the same as the gRNA AUG16 used in the experiments referenced in Figure 2C and 2D. (C) Relative Rluc luminescence with respect to Fluc luminescence was calculated to quantify the knockdown activity of WT Cas13 (left) and the dCas13 (right) variants using gRNAs targeting the start codon of Rluc as shown in B. (D) Relative Rluc mRNA abundance with respect to Fluc mRNA abundance was quantified by RT[qPCR under the indicated conditions. (E and G) Schematic of the luciferase reporter assay and gRNA design. gRNAs targeting various positions along the Rluc reporter mRNA were designed for both PspCas13b-NES and dPspCas13b-NES. A gRNA containing a nontargeting spacer sequence was used as a control. Fluc expression was used as an internal control. (F and H) Relative Rluc luminescence with respect to Fluc luminescence was calculated to quantify the knockdown activity of PspCas13b-NES (left) and dPspCas13b-NES (right) using the gRNAs shown in E and G. In C, D, F, and H, the mean (gray bar), s.d. (black line), and individual replicates (n = 3, black point) are shown. See also Figure S1.

To evaluate the potential of dCas13 to repress translation in mammalian cells, we used a dual reporter construct that expresses *Renilla* luciferase (Rluc) and firefly luciferase (Fluc). This setup allows one luciferase to act as a direct target for dCas13, whereas the other luciferase serves as an internal control. For each dCas13 protein, we designed a gRNA with a 30-nt-long spacer targeting the start codon of Rluc (Figures 1B and S1A). Transient coexpression of the Cas13/dCas13 protein, the respective gRNA, and the dual reporter allowed us to monitor the knockdown efficacy.

The dCas13 effectors reduced Rluc expression (Figure 1C). The impacts of dCas13 expression were generally correlated with those of the wild-type (WT) variants but were weaker (Figure 1C). Since among the constructs tested, dPspCas13b fused to NES (dPspCas13b-NES) showed the most robust effect, we focused mainly on this variant for subsequent assays (Figure 1C). While WT PspCas13b reduced Rluc mRNA, dPspCas13-induced Rluc repression was not associated with target RNA degradation (Figure 1D); hence, this repression was considered to stem from the net reduction in translation.

Given that dCas13 recruited to the start codon may physically obstruct the scanning 40S ribosome to impede translation initiation, we investigated the potency of dCas13 for inhibiting the elongating 80S ribosome. Testing of gRNAs targeting various positions along the reporter transcript (Figure 1E) revealed that dPspCas13-NES targeting the open reading frame (ORF) (gRNA ORF #1, ORF#2, and stop codon) led to weaker knockdown (Figure 1F). We also observed similar trends when dPspCas13b-NLS and dRfxCas13d-NLS were used for identical target sequences (Figure S1B-C). Notably, the defects in dCas13-mediated translational repression by the ORF-targeting gRNAs were not attributed to inactivity of the gRNAs, since they showed robust knockdown capacity in the presence of WT Cas13 (Figures 1F and S1B-C). These data indicated that dCas13 may not pose a strong enough blockade to halt the highly processive 80S, which can displace RNA-binding proteins from mRNA ^73^.

Considering the 40S scanning mechanism, we reasoned that, in addition to gRNAs covering the start codon, those targeting the 5′ untranslated region (UTR) may also induce translational repression. Thus, we designed gRNAs complementary to different positions in the 5′ UTR of the reporter (Figure 1G). Although reporter assays revealed a variation in the repressive capacity of dCas13for the sequences targeted by the tested gRNAs (Figure 1H), we observed translational repression for those gRNAs targeting the 5′ UTR. However, the gRNA targeting the start codon ensured the highest efficiency of translational repression. Experiments with dRfxCas13d-NLS led to conclusions similar to those reached for dPspCas13b-NES (Figure S1D). Therefore, we further optimized the gRNA contexts of the start codon in the dCas13-induced translational repression system.

### Construction of an optimal gRNA for translational repression

Given that gRNAs may have differential efficacy of target cleavage ^18, 72, 74–76^, we conducted a systematic survey to identify a gRNA design effective for translational inhibition. We tested the minimal length of the spacer region. Since the DR of the gRNA for PspCas13b is located at the 3′ end (Figure S1A), we retained the 3′ part of the spacer region and trimmed this region from the 5′ end (Figure 2A). gRNAs with short spacer regions, for example, those containing 16 and 17 nt, diminished the potency of translational repression, and 19- to 25-nt-long spacers provided the most efficient translational silencing (Figures 2B and S2A).

**Figure 2.**
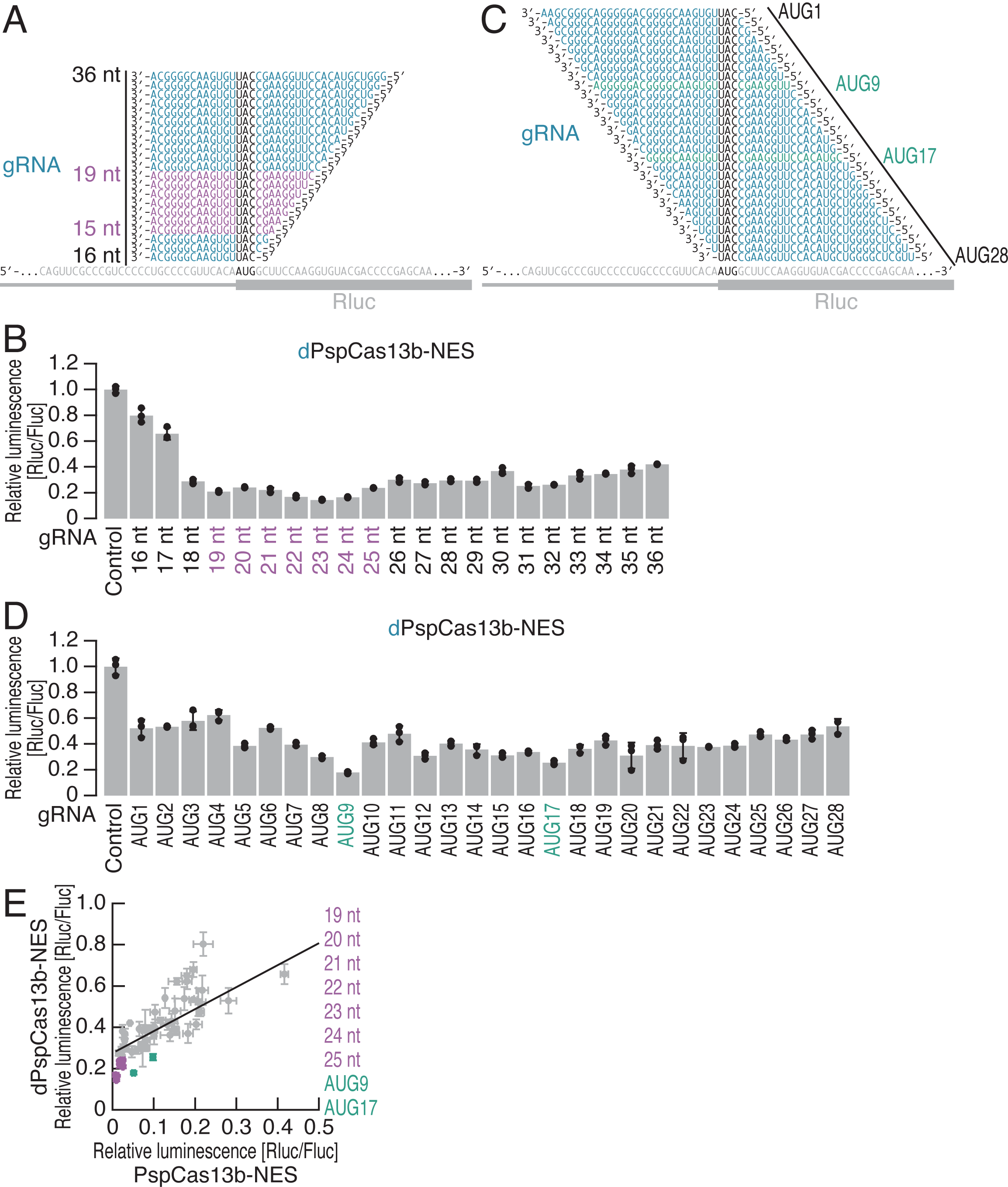
Optimal gRNA design for CRISPRδ. (A and C) Schematic of the gRNA design for targeting the start codon of Rluc reporter mRNA. The gRNAs with the highest efficacies are highlighted in purple (A) and green (C). (B and D) Relative Rluc luminescence with respect to Fluc luminescence was calculated to quantify the knockdown activity of dPspCas13b-NES using the gRNAs shown in A and C. The mean (gray bar), s.d. (black line), and individual replicates (n = 3, black point) are shown. The gRNAs with the highest efficacies are highlighted in purple (B) and green (D). (E) Comparison of the knockdown activity of PspCas13b-NES and dPspCas13b-NES across the gRNAs used in this study. The mean (gray point), s.d. (gray line), and regression line (black line) are shown. The gRNAs with the highest efficacies are highlighted in purple and green. See also Figure S2.

Since AUG masking showed potential as a valid strategy, we sought to determine whether the position of the anti-start codon within the spacer region affects the translational repression capacity of dCas13. For this purpose, we tiled 30-nt-long gRNAs along the start codon of Rluc (Figure 2C). Although we observed translational repression of Rluc with all gRNAs tested, central placement of the anti-start codon (*e.g.*, gRNA AUG9 and gRNA AUG17) in the spacer had a positive effect on translational repression (Figures 2D and S2B). A similar design principle was applicable to other dCas13 variants (dPspCas13b-NLS and dRfxCas13d-NLS) (Figure S2C-F) and gRNAs targeting Fluc (Figure S2G-H).

Generally, the knockdown potential of dPspCas13b-NES across the tested gRNAs scaled with that of the WT counterpart (Figure 2E), indicating that the efficacy of translational repression relies on the accessibility of the complex to the mRNA, as is the case for WT Cas13 ^72, 75, 76^. However, the gRNAs with the short spacer (19-25 nt) and those masking the AUG start codon in the center of the spacer (positions 9 and 17) exhibited an advantage for translational repression (Figure 2E).

Considering these observations, we termed our method of dCas13-mediated translational silencing CRISPRδ.

### High specificity of CRISPRδ

Given that WT Cas13 shows collateral RNase activity in mammalian cells ^21, 43–53^, we investigated the specificity of CRISPRδ. For this purpose, we performed ribosome profiling, a technique for deep sequencing of the ribosome-protected RNA fragments generated by RNase treatment ^77, 78^, and RNA-Seq from the same samples. We applied this approach in a cell line expressing enhanced GFP (EGFP) from a stable genomic integrant, with transient expression of Cas13 proteins and gRNAs. Here, we designed the gRNA to target EGFP (Figure 3A) in accordance with the configuration of the effective gRNA in the luciferase-based assay (Figure 2). Since the stochastic expression of Cas13 proteins in cells hampers quantitative analysis, we sorted cells by flow cytometry to ensure the expression of mCherry-tagged Cas13 in the cells. Indeed, we observed that the cells with higher expression of mCherry-tagged Cas13 (WT or catalytically dead) showed more potent repression of EGFP (Figure S3A and S3B).

**Figure 3.**
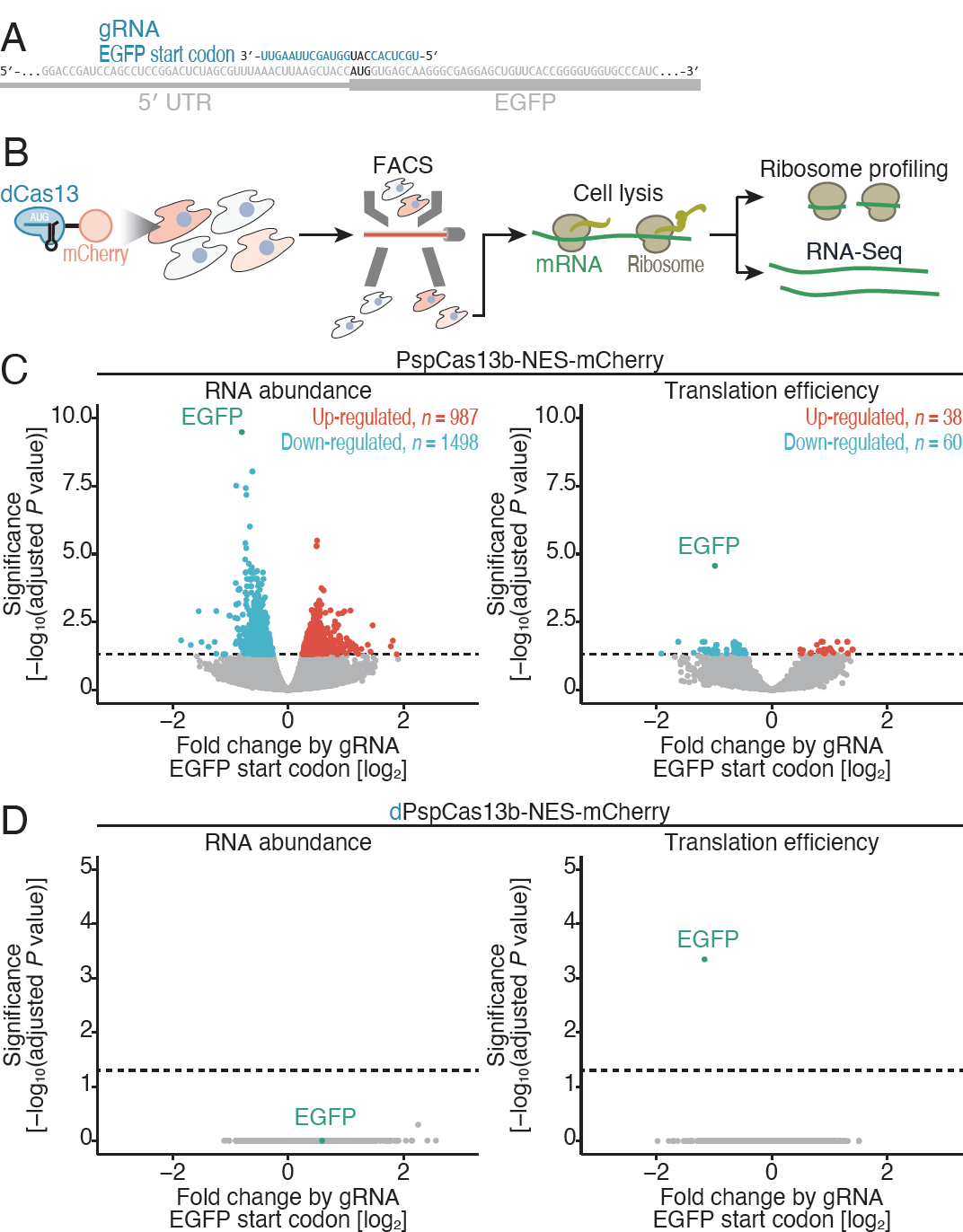
Ribosome profiling and RNA-Seq reveal the high specificity of CRISPRδ for gene knockdown. (A) Schematic of the gRNA design for targeting the start codon of the EGFP reporter mRNA. (B) Schematic of experimental procedures for ribosome profiling and RNA-Seq. PspCas13b variant expression was ensured by cell sorting. (C) Volcano plots of significance (*P* value adjusted by the Benjamini–Hochberg method) and fold change in the RNA abundance (left) and translation efficiency (right) by expression of the gRNA targeting the EGFP start codon and WT PspCas13b-NES. mRNAs with significant alterations (adjusted *P* value < 0.05) are highlighted. (D) Same as C but for dPspCas13b-NES. See also Figure S3.

Combined fluorescence-activated cell sorting (FACS) and ribosome profiling/RNA-Seq were performed to evaluate the effects of the Cas13 variants on gene expression in a genome-wide manner (Figure 3B). The hallmarks of the read features, such as the peak footprint lengths of 29-30 nt and 21-22 nt (Figure S3C) ^79, 80^ and the 3-nt periodicity along the ORF (Figure S3D), validated the quality of the ribosome profiling in the cells collected by FACS. As reported in earlier studies ^21, 43–53^, the collateral activity of WT PspCas13b-NES induced changes in the expression of nontarget mRNAs (987 upregulated transcripts and 1498 downregulated transcripts), in addition to the intended reduction in EGFP expression (Figure 3C left). In contrast, dPspCas13b-NES did not lead to any transcriptomic changes (including the expression of EGFP) (Figure 3D left). However, in terms of translation efficiency—that is, the over- or underrepresentation of ribosome footprints with respect to the changes in RNA abundance—dPspCas13b-NES enabled suppression of EGFP expression (Figure 3D right). Importantly, this suppression was highly specific for this gRNA-targeted transcript; we found no significant alterations in mRNA expression across the transcriptome (Figure 3D right).

Notably, even WT PspCas13b-NES may induce translational alterations in the target EGFP transcript and other transcripts (Figure 3C right). Translational repression of EGFP probably occurs because the recruitment of the WT variant also results in steric hindrance of scanning ribosomes before transcript degradation. In addition, ribotoxic stress ^81, 82^ due to partial cleavage of ribosomal RNA (rRNA) by WT Cas13 ^51^ may elicit changes in the translation of nontarget mRNAs. Again, we did not find that dPspCas13b-NES exerted such a nonspecific effect on translation (Figure 3D right).

Taking these data together, we concluded that the CRISPRδ method can be used for gene silencing with ultrahigh specificity.

### CRISPR**δ** represses cap-independent translation

Although most mRNAs utilize the cap-dependent process to initiate translation in eukaryotic cells ^83^, a subset of mRNAs exploit cap-independent mechanisms such as internal ribosome entry sites (IRES)-mediated translation ^84, 85^. To explore the ability of CRISPRδ to repress IRES-mediated protein synthesis, we used an Rluc reporter fused to the hepatitis C virus (HCV) IRES ^86, 87^, which directly recruits the 40S ribosome to the AUG codon and bypasses the cap-binding protein eIF4E, scaffold protein eIF4G, and RNA-binding protein eIF4A. Indeed, hippuristanol, a eIF4A inhibitor ^88, 89^, treatment confirmed that our reporter was translated independently of the protein (Figure S4A).

dPspCas13b-NES repressed translation driven by the HCV IRES (Figure 4A and 4B). This effect was specific for the gRNA targeting the AUG codon of the HCV IRES. The downstream AUG codon, which was initially used as the start codon of Rluc (Figure 4A), could not serve as a valid target (Figure 4B), corroborating the idea that dCas13 is not potent for blocking translation elongation.

**Figure 4.**
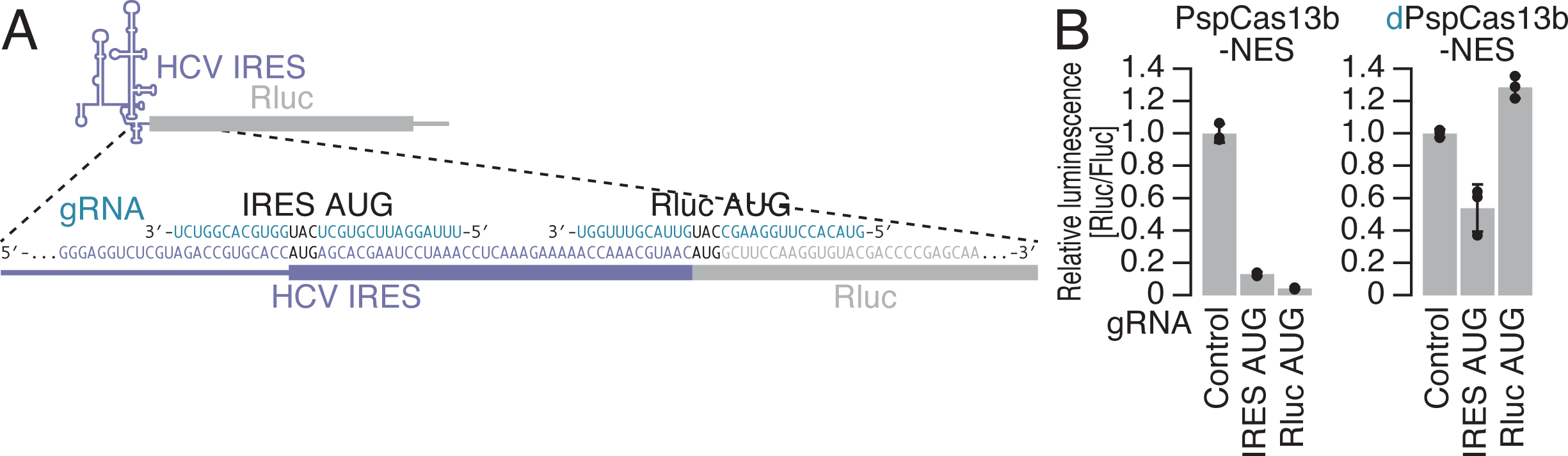
CRISPRδ represses IRES-driven translation. (A) Schematic of the gRNA design for targeting the start codon of Rluc reporter mRNA containing the HCV IRES. (B) Relative Rluc luminescence with respect to Fluc luminescence was calculated to quantify the knockdown activity of dPspCas13b-NES using the gRNAs shown in A. The mean (gray bar), s.d. (black line), and individual replicates (n = 3, black point) are shown. See also Figure S4.

### Enhanced CRISPR**δ** system

Despite the high specificity of CRISPRδ with dCas13, the efficacy was modest compared to that with WT Cas13. Whereas WT Cas13 can catalytically degrade target mRNAs in multiple rounds of reactions, dCas13 is required to associate with the target mRNA for a long enough duration to suppress translation and thus needs to be stoichiometrically abundant. To facilitate the effect of dCas13 on translational repression, we tested multiple gRNAs targeting a single mRNA; two different gRNAs were used to increase the duration of dCas13-mRNA association (Figure S5A). This strategy enabled further downregulation of target mRNA translation by gRNAs (Figure S5A), which originally showed modest translational repression activity.

We also tested the fusion of a translational repressor to dCas13. This option has been successfully used to boost dCas9-mediated transcriptional repression in CRISPR interference (CRISPRi) ^90–93^. Considering the steric hindrance effect of dCas13 on scanning 40S ribosomes, another mechanism of translational repression by the fused protein may further benefit translational repression (Figure 5A). Here, we tested several translational suppressor proteins, including programmed cell death 4 (PDCD4) ^94–97^, eIF4E-transporter protein (4E-T) ^98–100^, eIF6 ^101, 102^, 14-3-3σ ^103^, poly(A)-binding protein-interacting protein 2 (PAIP2) ^104–109^, Pelota (PELO) ^110–114^, and 4EHP ^115–120^, to dPspCas13b-NES targeting the start codon of the reporter and found that 4EHP exerted the most significant effects (Figure S5B). The recruitment of 4EHP-fused dPspCas13b-NES not only to the start codon but also to the 5′ UTR outperformed that of the nonfused variant (Figure 5B). Moreover, this fusion system allowed the inhibition of protein synthesis with a gRNA targeting the 3′ UTR, which originally could not impact translation with nonfused dCas13 (Figures 1F and 5B), as shown by artificial tethering of this protein ^118, 120^

**Figure 5.**
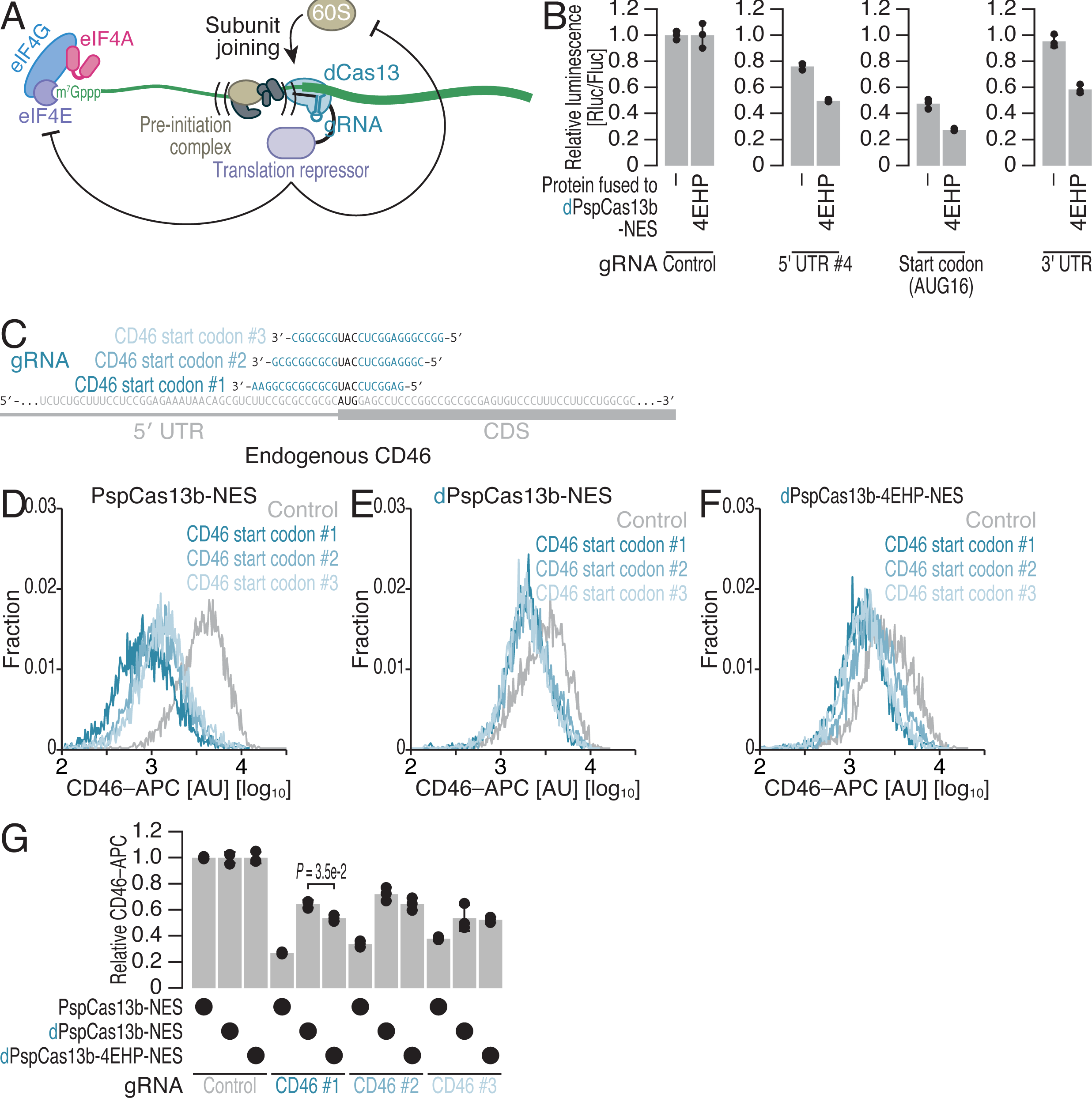
Fusion of translational repressors to dPspCas13b enhances knockdown efficacy. (A) Schematic showing the enhancement of dCas13-induced repression of mRNA translation by fusion of a translational repressor. Fused translational repressors provide an alternative mode of suppression. (B) Relative Rluc luminescence with respect to Fluc luminescence was calculated to quantify the knockdown activity of dPspCas13b-NES and dPspCas13b-4EHP-NES using the gRNAs shown in Figure 1E. The mean (gray bar), s.d. (black line), and individual replicates (n = 3, black point) are shown. (C) Schematic of the gRNA design for targeting the start codon of CD46 mRNA with PspCas13b-NES/dPspCas13b-NES/dPspCas13b-4EHP-NES. (D-F) Representative distribution of APC-labeled CD46 with the expression of the indicated PspCas13 variants and gRNAs. Typically, 4,000-5,000 cells, which passed the expression threshold of mCherry fused to PspCas13 variants, were considered. (G) Quantification of CD46 expression by FACS. The average values for the distributions (D-F) were used. The mean (gray bar), s.d. (black line), and individual replicates (n = 3, black point) are shown. Significance was determined by Student’s t test (two-tailed). See also Figure S5.

### CRISPR**δ** silences endogenous mRNA translation

To test the potency of the enhanced CRISPRδ system, we applied it to endogenous cellular mRNAs. Here, we designed three gRNAs targeting the CD46 start codons (Figure 5C) for the PspCas13b variants and measured the abundance of CD46 protein on the cell surface with flow cytometry. In addition to WT PspCas13b-NES (Figure 5D), dPspCas13b-NES suppressed CD46 protein expression (Figure 5E). The efficacy was further augmented by 4EHP fusion (Figure 5F and 5G, gRNA CD46 #1).

Taken together, these data established CRISPRδ as a potent and highly specific tool for gene silencing through translational repression, independent of RNA degradation.

## Discussion

To overcome the collateral activity and toxicity stemming from the use of Cas13 as a gene knockdown tool ^21, 43–53^, the use of engineered Cas13 proteins ^53^, the controlled expression of the Cas 13 protein ^50^, and the use of Cas13 variants with lower collateral activity ^48^ have been proposed. Despite the promising results shown in the reports, these approaches could not completely prevent RNase activity toward bystander RNA species. In contrast, the CRISPRδ system presented in this study does not rely at all on RNA degradation and thus allows gene knockdown without the risk of induced collateral activity. Instead, CRISPRδ harnesses the high specificity of targeting by the dCas13-gRNA complex and the tight interaction of this complex with the target, resulting in steric hindrance of scanning ribosomes. This may be a sensitive means for general translational control since similar mechanisms are employed by natural translational regulators ^121, 122^.

Recent reports have suggested that dCas13 could be a useful tool to modulate translation. In bacteria, targeting the Shine-Dalgarno (SD) sequence upstream of the start codon by dLwaCas13a or dRfxCas13d prevents hybridization of the anti-SD sequence in 16S rRNA and results in translational repression *in vitro* ^123^ and *in vivo* ^124^. The mechanism of inhibition in the bacterial studies and this work differed due to the different principles of translation initiation in prokaryotes (internal recruitment of the small ribosome subunit via the SD/anti-SD interaction) and eukaryotes (scanning by the preinitiation complex) ^83^. However, all the works shared the strategy of targeting a process before the formation of elongating ribosomes. Although those earlier studies lacked genome-wide investigation, this strategy may also have high specificity for gene knockdown in bacteria, as highlighted in this work. In addition to inducing translational repression, the dCas13 approach may be able to activate protein synthesis. Targeting IF3-fused dRfxCas13d to the 5′ UTR may augment translation in bacteria ^124^. In mammals, conjugation of the SINEB2 repeat element, which activates protein synthesis from hybridized mRNA ^125^, to the gRNA of dRfxCa13d enhances translation ^126^. In further studies, the applications of dCas13 will be expanded for translational control in a wide range of biological contexts across diverse organisms.

We expect translational repression mediated by CRISPRδ to have several advantages over preexisting methods. A wide array of studies have reported small but functional micropeptides encoded by long noncoding RNAs ^127, 128^, which were originally expected to contain no protein-coding regions. Moreover, mRNAs have been reported to have protein coding-independent functions ^129–131^. Thus, our method, which suppresses protein synthesis but leaves the target RNA intact, may be useful in delineating the roles of the RNA in protein production or other roles. Moreover, CRISPRδ is expected to circumvent the effects of the genetic compensation mechanism elicited by gene knockout by nonsense mutation ^132, 133^, allowing the straightforward interpretation of the gene expression-phenotype interaction. CRISPRδ represents a valuable step toward the development of a toolbox for reliable gene knockdown and for a new clarification of gene function.

## Acknowledgments

We are grateful to all the members of the Iwasaki laboratory for constructive discussions and technical help. We also thank the Support Unit for Bio-Material Analysis, RIKEN CBS Research Resources Division, for Sanger sequencing and FACS analysis. Computation was supported by the HOKUSAI SailingShip supercomputer facility at RIKEN. We also thank Dr. Junichi Tanaka for sharing hippuristanol with us. The pCAGEN vector was a kind gift from Dr. Yukihide Tomari. S.I. was supported by the Ministry of Education, Culture, Sports, Science and Technology (MEXT) [a Grant-in-Aid for Transformative Research Areas (B) “Parametric Translation”, JP20H05784]; the Japan Society for the Promotion of Science (JSPS) [a Grant-in-Aid for Scientific Research (B), JP23H02415]; the Japan Agency for Medical Research and Development (AMED) (AMED-CREST, JP23gm1410001); and RIKEN (Pioneering Projects “Biology of Intracellular Environments”). H.T. was supported by AMED (AMED-CREST, JP23gm1410001). Y.S. was supported by JSPS [a Grant-in-Aid for Early-Career Scientists, JP21K15023; Grant-in-Aid for Scientific Research (C), JP23K05648]; MEXT [a Grant-in-Aid for Transformative Research Areas (A) “Multifaceted Proteins”, JP21H05734, JP23H04268]; and RIKEN (Special Postdoctoral Researchers and Incentive Research Projects). Y.S. was a recipient of the RIKEN Special Postdoctoral Researchers Program. A.A. was an International Program Associate of RIKEN.

## Author contributions

Conceptualization: A.A., H.T., Y.S., and S.I.;

Methodology: A.A., Y.S., and S.I.;

Formal Analysis: A.A.;

Investigation: A.A.;

Writing – Original Draft: A.A., Y.S., and S.I.;

Writing – Review & Editing: A.A., H.T., Y.S., and S.I.;

Visualization: A.A. and S.I.;

Supervision: Y.S. and S.I.;

Funding Acquisition: H.T., Y.S., and S.I.

## Declaration of interests

The authors declare that they have no competing interests.

## Experimental procedures

### Cell cultures

The human embryonic kidney (HEK) 293 [American Type Culture Collection (ATCC), CRL-1573] line was cultured in high-glucose DMEM containing GlutaMAX supplement (Thermo Fisher Scientific) and 10% fetal bovine serum (FBS; Sigma–Aldrich) at 37°C with 5% CO_2_. Cell cultures were routinely tested for contamination with *Mycoplasma spp.* (e-Myco VALiD Mycoplasma PCR Detection Kit, iNtRON Biotechnology).

To establish a stable cell line expressing EGFP, HEK293 Flp-In T-Rex cells (Thermo Fisher Scientific, R78007) were transfected with pcDNA5/FRT/TO-GFP (see below) and pOG44 (Thermo Fisher Scientific) using X-tremeGENE9 (Roche) and cultured in the presence of blasticidin S (InvivoGen) and hygromycin B (InvivoGen).

### Constructs for Cas13 expression

#### pXR-LwaCas13a-NLS, LwaCas13a-NES, dLwaCas13a-NLS, and dLwaCas13a-NES

The DNA fragment encoding dLwaCas13a was PCR-amplified using pC035-dLwaCas13a-msfGFP (a gift from Feng Zhang; Addgene plasmid #91925; http://n2t.net/addgene:91925; RRID: Addgene_91925) ^17^ as a template and inserted into pXR002: EF1a-dCasRx-2A-EGFP (a gift from Patrick Hsu; Addgene plasmid #109050; http://n2t.net/addgene:109050; RRID: Addgene_109050) ^19^ between the C-terminal and N-terminal NLSs in the plasmid. To generate the plasmid that contained the NES, the DNA fragment encoding dLwaCas13a fused to a C-terminal NES sequence was PCR–amplified and inserted into pXR002: EF1a-dCasRx-2A-EGFP, eliminating the NLSs. To convert dLwaCas13a to the catalytically active WT LwaCas13a, the A474R and A1046R substitutions were generated by site-directed mutagenesis.

#### pXR-PguCas13-NLS, PguCas13-NES, dPguCas13-NLS, and dPguCas13-NES

The DNA fragment encoding PguCas13b was PCR-amplified, using pC0045-EF1a-PguCas13b-NES-HIV (a gift from Feng Zhang; Addgene plasmid #103861; http://n2t.net/addgene:103861; RRID: Addgene_103861) ^18^ as a template and inserted into pXR002: EF1a-dCasRx-2A-EGFP ^19^ between the C-terminal and N-terminal NLSs in the plasmid. To generate the plasmid that contained the NES, the DNA fragment encoding PguCas13b fused to a C-terminal NES sequence was PCR-amplified and inserted into pXR002: EF1a-dCasRx-2A-EGFP, eliminating the NLSs. To convert PguCas13b to catalytically inactive dPguCas13b mutant, the H151A and H1121A substitutions were generated by site-directed mutagenesis.

#### pXR-RfxCas13d-NLS, RfxCas13d-NES, dRfxCas13d-NLS, and dRfxCas13d-NES

pXR-dRfxCas13d-NLS was repotered in the earlier study (originally named as pXR002: EF1a-dCasRx-2A-EGFP) ^19^. To generate the plasmid that contained the NES, the DNA fragment encoding dRfxCas13d fused to a C-terminal NES sequence was PCR-amplified and inserted into pXR002: EF1a-dCasRx-2A-EGFP, eliminating the NLSs. To convert dRfxCas13d to catalytically active WT RfxCas13d, the A239R, A244H, A858R, and A863H substitutions were generated by site-directed mutagenesis.

#### pXR-PspCas13b-NLS, PspCas13b-NES, dPspCas13b-NLS, and dPspCas13b-NES

The DNA fragment encoding dPspCas13b was PCR-amplified, using pC0049-EF1a-dPSPCas13b-NES-HIV, H133A/H1058A (a gift from Feng Zhang; Addgene plasmid #103865; http://n2t.net/addgene:103865; RRID: Addgene_103865) ^18^ as a template and inserted into pXR002: EF1a-dCasRx-2A-EGFP ^19^ the C-terminal and N-terminal NLSs in the plasmid. To generate the plasmid that contained the NES, the DNA fragment encoding dPspCas13b fused to a C-terminal NES sequence was PCR-amplified and inserted into pXR002: EF1a-dCasRx-2A-EGFP, eliminating the NLSs. To convert dPspCas13b to catalytically active WT PspCas13b, the A133H and A1058H substitutions were generated by site-directed mutagenesis.

#### pXR-dPspCas13b-PDCD4-NES, 4E-T-NES, eIF6-NES, 14-3-3σ-NES, PAIP2-NES, PELO-NES, and 4EHP-NES

The DNA fragments encoding the translational repressor proteins (PDCD4, 4E-T, eIF6, 14-3-3σ, PAIP2, PELO, or 4EHP) were synthesized (Eurofins Genomics), used as templates for PCR amplification, and fused to an XTEN10 linker ^134, 135^. The amplified fragments were then inserted between the dPspCas13b and NES sequences in pXR-dPspCas13b-NES.

#### pCAGEN-PspCas13b-NES-mCherry, dPspCas13b-NES-mCherry, and PspCas13b-4EHP-NES-mCherry

The DNA fragments encoding PspCas13b-NES, dPspCas13b-NES, and dPspCas13b-4EHP-NES were PCR-amplified, using pXR-PspCas13b-NES, pXR-dPspCas13b-NES, and pXR-dPspCas13b-4EHP-NES, respectively, as a template and inserted into the pCAGEN vector (a gift from Dr. Yukihide Tomari at The University of Tokyo) ^136^, along with a PCR-amplified DNA fragment encoding mCherry.

### Constructs for gRNA expression

To construct the gRNA-expressing plasmids, the DR corresponding to each Cas13 ortholog, along with the respective spacer sequence, was inserted into the pC016-LwCas13a guide expression backbone with U6 promoter (a gift from Feng Zhang; Addgene plasmid #91906; http://n2t.net/addgene:91906; RRID: Addgene_91906). The gRNA expression plasmids are listed in Table S1.

### Constructs for reporter expression

#### psiCHECK2-PTGES3

psiCHECK2-PTGES3, which contains the Rluc ORF fused to the PTGE3 5′ UTR, was reported previously ^122^.

#### psiCHECK2-HCV IRES

psiCHECK2-HCV IRES, which contains the Rluc ORF fused to the HCV IRES, was reported previously ^122^.

#### pCDNA5/FRT/TO-GFP

The PCR-amplified fragment encoding EGFP was inserted between the Hind III and BamH I sites in pcDNA5/FRT/TO (Thermo Fisher Scientific, V6520-20).

### Luciferase reporter assay

#### Rluc and Fluc dual assay

Approximately 1 × 10^5^ HEK293 cells were seeded in flat-bottom 24-well plates (Thermo Fisher Scientific or FALCON) and incubated overnight. The following day, the cells were cotransfected with the Cas13 variant-expressing plasmid (pXR series, 150 ng), the gRNA-expressing plasmid (pC016 series, 300 ng), and the luciferase reporter plasmid (psiCHECK2-PTGES3, 5 ng; psiCHECK2-HCV IRES, 10 ng) using Lipofectamine 3000 (Thermo Fisher Scientific) according to the manufacturer’s instructions. Forty-eight hours post-transfection, cells were lysed with 100 μl of Passive Lysis Buffer (Promega). Then, 10 μl of lysate from each well was transferred to a 96-well flat-bottom white assay plate (Corning), and luminescence was measured by the Dual-Luciferase Reporter Assay System (Promega) and GloMax Navigator System (Promega).

For the confirmation of IRES-mediated translation initiation, 10 ng of psiCHECK2-PTGES3 or psiCHECK2-HCV IRES was used. Hippuristanol (1 μM with 0.1% DMSO as a stock solvent) (a gift from Dr. Junichi Tanaka at University of the Ryukyus) or DMSO (0.1%) was added to the cells 10 h before lysis.

#### Data analysis

The average of the background luminescence signals (Rluc, and Fluc) from triplicate wells of nontransfected cells was subtracted from the luminescence values of the experimental wells. The luminescence values were normalized to the average of the corresponding triplicate control wells.

### RT⃩qPCR

Approximately 4 × 10^5^ cells were seeded in flat-bottom 6-well plates (Thermo Fisher Scientific) and incubated overnight. The following day, the cells were cotransfected with the Cas13 variant-expressing plasmid (pXR series, 600 ng), the gRNA-expressing plasmid (pC016 series, 1200 ng), and the luciferase reporter plasmid (psiCHECK2-PTGES3, 20 ng) using Lipofectamine 3000 (Thermo Fisher Scientific) according to the manufacturer’s instructions. RNA was harvested 48 h post-transfection using TRIzol reagent (Thermo Fisher Scientific) and Direct-zol RNA MicroPrep Kit (Zymo Research) according to the manufacturers’ instructions. Then, RNA was treated with DNase (TURBO DNA-*free* Kit, Thermo Fisher Scientific), purified with RNAClean XP beads (Beckman Coulter), and reverse-transcribed using ReverTra Ace qPCR RT Master Mix (TOYOBO). qPCR was performed on a thermal cycler (TaKaRa) using TB Green Premix Ex TaqTM II (TaKaRa) with the following primers: Rluc, 5′-TCGTCCATGCTGAGAGTGTC-3′ and 5′-CTAACCTCGCCCTTCTCCTT-3′; Fluc, 5′-TTCGCTAAGAGCACCCTGAT-3′ and 5′-GTAATCAGAATGGCGCTGGT-3′.

### FACS

#### Assessment of EGFP expression

Approximately 4 × 10^6^ HEK293 Flp-In T-Rex cells with EGFP integrants were seeded in 15-cm dishes (Thermo Fisher Scientific) and incubated overnight. The following day, the cells were cotransfected with the Cas13 variant-expressing plasmid (pCAGEN-PspCas13b-NES-mCherry or pCAGEN-dPspCas13b-NES-mCherry, 8 μg) and the gRNA-expressing plasmid (pC016-gPsp-Control or pC016-gPsp-EGFP start codon, 12 μg) using TransIT-293 (Mirus) according to the manufacturer’s instructions. The cells were then incubated for 48 h prior to FACS. To induce EGFP expression, the cells were treated with tetracycline (1 μg/ml) for 24 h prior to FACS.

The cells were washed with PBS (Nacalai Tesque) and trypsinized (Thermo Fisher Scientific). Then, the cells were resuspended in high-glucose DMEM containing GlutaMAX supplement (Thermo Fisher Scientific) and 10% FBS (Sigma–Aldrich) and were centrifuged at 800 × *g* for 3 min. After the supernatant was removed, the cells were washed with PBS and centrifuged again at 800 × *g* for 3 min. The cell pellet was resuspended in 1.5 ml of PBS. After passing through a 35-μm mesh filter (FALCON), cells were analyzed in FACSAria II Special Order system (BD) to measure EGFP and mCherry expression. Data from 1 × 10^4^ cells were used for analysis.

#### Assessment of CD46 expression

Approximately 4 × 10^5^ cells were seeded in flat-bottom 6-well plates (Thermo Fisher Scientific) and incubated overnight. The following day, the cells were cotransfected with the Cas13 variant-expressing plasmid (pCAGEN-PspCas13b-NES-mCherry, pCAGEN-dPspCas13b-NES-mCherry, or pCAGEN-dPspCas13b-4EHP-NES-mCherry, 800 ng) and the gRNA-expressing plasmid (pC016-gPsp-Control, pC016-gPsp-CD46 #1, pC016-gPsp-CD46 #2, or pC016-gPsp-CD46 #3, 1200 ng) using TransIT 293 (Mirus) according to the manufacturer’s instructions. The cells were then incubated for 48 h prior to FACS.

The cells were washed with PBS (Nacalai Tesque) and trypsinized (Thermo Fisher Scientific). Then, the cells were resuspended in high-glucose DMEM containing GlutaMAX supplement (Thermo Fisher Scientific) and 10% FBS (Sigma–Aldrich) and centrifuged at 800 × *g* for 3 min. After the supernatant was removed, the cells were washed with FACS Buffer [2% FBS and 0.1% NaN_3_ (Nacalai Tesque) in PBS (Nacalai Tesque)] and centrifuged at 800 × *g* for 3 min. The cells were resuspended in 10 μl of FACS Buffer and incubated with 1 μl of allophycocyanin (APC)-conjugated anti-CD46 antibody (BioLegend, clone TRA-2–10, 352405) for 30 min at 4°C with rotation. Then, the cells were washed twice with FACS Buffer and resuspended in 1 ml of FACS Buffer. After passing through a 35-μm mesh filter (FALCON), cells were analyzed in FACSAria II Special Order system (BD) to measure APC signals and mCherry expression. The background signal for mCherry was measured in cells not transfected with an mCherry-fused Cas13 variant expression plasmid and used for the identification of mCherry-positive cells (a value of 10^3^ or higher).

### Ribosome profiling and RNA-Seq

#### Library preparation

The ribosome profiling library was prepared as described previously ^137^ with modifications. Cell seeding, transfection, and sorting were prepared as described in “*Assessment of EGFP expression*”. Cycloheximide (100 μg/ml, Sigma–Aldrich) was added to PBS, trypsin solution, and DMEM to halt ribosome movement along mRNAs in the harvested cells. Then, the cells were subjected to FACS as described in “*Assessment of EGFP expression*”. As a control for cell sorting, we used cells that were not transfected with the plasmid for mCherry-fused Cas13 variant expression and collected cells with a value of 10^2^ or higher as mCherry-positive cells. The sorted 1 × 10^6^ cells were pelleted by centrifugation at 800 × *g* for 3 min and subsequently lysed in lysis buffer (20 mM Tris-HCl pH 7.5, 150 mM NaCl, 5 mM MgCl_2_ 1 mM DTT, 1% Triton X-100, 100 μg/ml cycloheximide, and 100 μg/ml chloramphenicol). The lysates were treated with Turbo DNase (Thermo Fisher Scientific) for 10 min at 4°C and then clarified by removal of cellular debris by centrifugation at 20,000[×[*g* for 10 min at 4°C. The total RNA concentration in each lysate was measured using a Qubit RNA BR Assay Kit (Thermo Fisher Scientific). Lysates containing 3 μg of total RNA were treated with 20 U of RNase I (Lucigen) for 45 min at 25°C and then subjected to a sucrose cushion ultracentrifugation at 100,000 rpm for 1 h at 4°C with Optima MAX-TL ultracentrifuge and TLA-110 rotor (Beckman Coulter). The RNA fragments in the resulting pellet were recovered using TRIzol reagent (Thermo Fisher Scientific) and Direct-zol RNA MicroPrep Kit (Zymo Research) and subjected to denaturing polyacrylamide gel electrophoresis (PAGE). RNA fragments with lengths ranging between 17 and 34 nt were excised from the gel. After purification, the RNA fragments were dephosphorylated by incubation with T4 polynucleotide kinase (New England Biolabs) for 1 h at 37°C and ligated to preadenylated linker oligonucleotides (5′-App NNNNNIIIIIAGATCGGAAGAGCACACGTCTGAA-ddC-3′, where App indicates preadenylation, ddC indicates 2′,3′-dideoxycytidine, Ns indicates a random sequence for the unique molecular index, and Is indicates the index sequence used for multiplexing) by incubation with T4 RNA Ligase 2, truncated KQ (New England Biolabs) for 3 h at 22°C in 17.5% PEG-8000. The linker-ligated RNA was subjected to denaturing PAGE and gel extraction. rRNA was removed from the purified RNA using Ribo-Zero Gold rRNA Removal Kit accompanied by TruSeq Stranded Total RNA Kit (Illumina). RNA was then reverse-transcribed for 30 min at 50°C using ProtoScript II (New England Biolabs) with a primer (5′-Phos-NNAGATCGGAAGAGCGTCGTGTAGGGAAAGAG-iSp18-GTGACTGGA GTTCAGACGTGTGCTC-3′, where Phos indicates 5′ monophosphate and iSp18 indicates an 18-atom hexaethylene glycol spacer). The template RNA was hydrolyzed with 1 M NaOH and subjected to denaturing PAGE and gel extraction. The purified cDNA was then circularized using CircLigaseII ssDNA ligase (Epicentre) for 1 h at 60°C and was subsequently PCR amplified using the primers 5′-AATGATACGGCGACCACCGAGATCTACACTCTTTCCCTACACGACGCTC-3′ and 5′-CAAGCAGAAGACGGCATACGAGATJJJJJJGTGACTGGAGTTCAGACGTGTG-3′(where Js indicates the 6-mer index sequence for Illumina sequencing).

For RNA-Seq library preparation, total RNA was extracted from the same lysate used for ribosome profiling using TRIzol LS reagent (Thermo Fisher Scientific) and Direct-zol RNA MiniPrep Kit (Zymo Research). The library was prepared using a TruSeq RNA Library Preparation Kit v2 (Illumina) according to the manufacturer’s instructions.

DNA libraries were sequenced on the HiSeq X (Illumina) platform with the paired-end 150-nt option.

#### Data analysis

Sequencing data were analyzed as described previously ^138, 139^. To perform read quality filtering and adapter trimming, we used fastp (ver. 0.21.0) ^140^. STAR (ver. 2.7.0a) ^141^ was used to remove reads aligned to noncoding RNAs. The remaining reads were mapped to the hg38 human genome by STAR and annotated with the GENCODE Human Release 32 reference obtained via the UCSC Genome Browser (https://genome.ucsc.edu/index.html).

We determined the offset of the A site in the reads according to the metagene analysis conducted with a custom script (https://github.com/ingolia-lab/RiboSeq) to be 15 for 25-33-nt reads. For RNA-Seq, we used an offset of 15 for all mRNA fragments. To calculate the changes in RNA abundances and translation efficiencies, which were calculated as ribosome profiling counts normalized to RNA-Seq counts, we used the DESeq2 package ^142^. We calculated the significance with the likelihood ratio test in a generalized linear model. The reads that corresponded to the first five codons and the last five codons in the ORF were excluded from our analyses. All custom-made scripts used in this study are available upon request.

### Accession numbers

The ribosome profiling and RNA-Seq data (GSE232383) obtained in this study were deposited in the National Center for Biotechnology Information (NCBI) database.

## Figure and table legends

**Figure S1.**
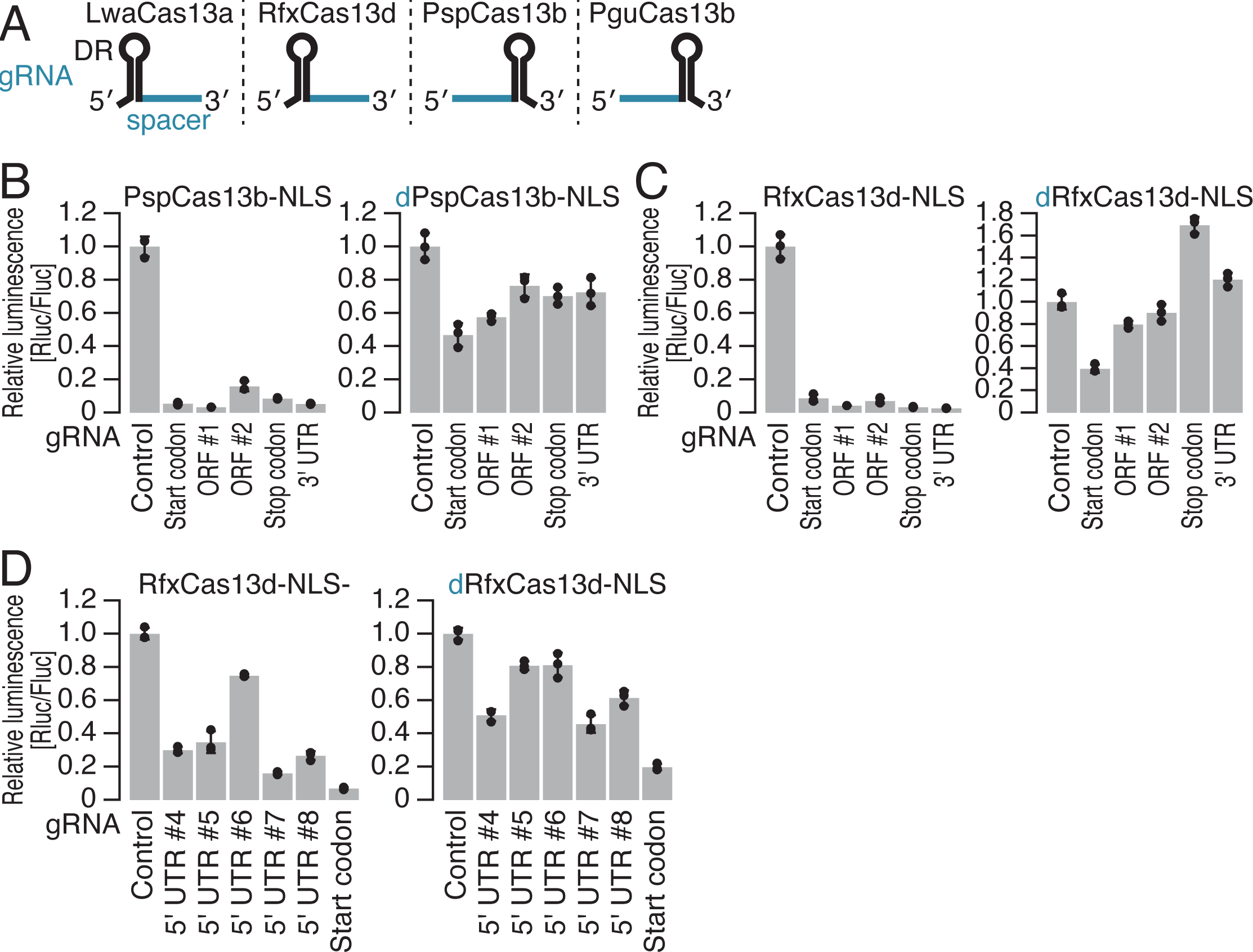
Characterization of Cas13 protein potency for translational repression, related to Figure 1. (A) Schematic of the gRNA architecture for each Cas13 protein. (B) Relative Rluc luminescence with respect to Fluc luminescence was calculated to quantify the knockdown activity of PspCas13b-NLS (left) and dPspCas13b-NLS (right) using the gRNAs shown in Figure 1E. (C) Relative Rluc luminescence with respect to Fluc luminescence was calculated to quantify the knockdown activity of RfxCas13d-NLS (left) and dRfxCas13d-NLS (right) using the gRNAs shown in Figure 1E. (D) Relative Rluc luminescence with respect to Fluc luminescence was calculated to quantify the knockdown activity of RfxCas13d-NLS (left) and dRfxCas13d-NLS (right) using the gRNAs shown in Figure 1G. In B-D, the mean (gray bar), s.d. (black line), and individual replicates (n = 3, black point) are shown.

**Figure S2.**
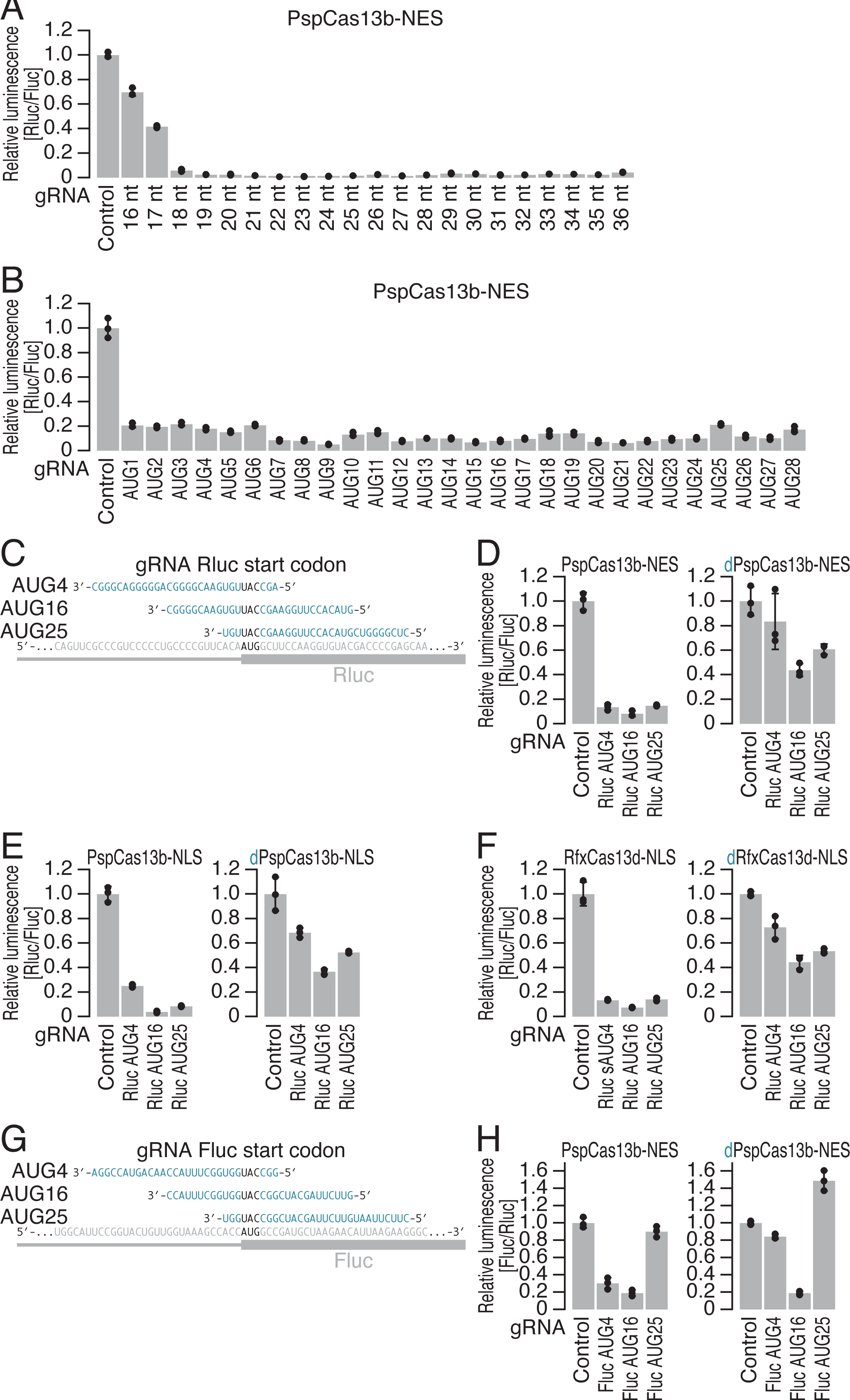
Characterization of gRNA potency for translational repression, related to Figure 2. (A and B) Relative Rluc luminescence with respect to Fluc luminescence was calculated to quantify the knockdown activity of PspCas13b-NES using the gRNAs shown in Figure 2A and C. The mean (gray bar), s.d. (black line), and individual replicates (n = 3, black point) are shown. (C and G) Schematic of the gRNA design for targeting the start codon of Rluc reporter mRNA (C) and Fluc reporter mRNA (G). (D) Relative Rluc luminescence with respect to Fluc luminescence was calculated to quantify the knockdown activity of PspCas13b-NES (left) and dPspCas13b-NES (right) using the gRNAs shown in C. (E) Relative Rluc luminescence with respect to Fluc luminescence was calculated to quantify the knockdown activity of PspCas13b-NLS (left) and dPspCas13b-NLS (right) using the gRNAs shown in C. (F) Relative Rluc luminescence with respect to Fluc luminescence was calculated to quantify the knockdown activity of RfxCas13d-NLS (left) and dRfxCas13d-NLS (right) using the gRNAs shown in C. (H) Relative Fluc luminescence with respect to Rluc luminescence was calculated to quantify the knockdown activity of PspCas13b-NES (left) and dPspCas13b-NES (right) using the gRNAs shown in G. In A, B, D-F, and H, the mean (gray bar), s.d. (black line), and individual replicates (n = 3, black point) are shown.

**Figure S3.**
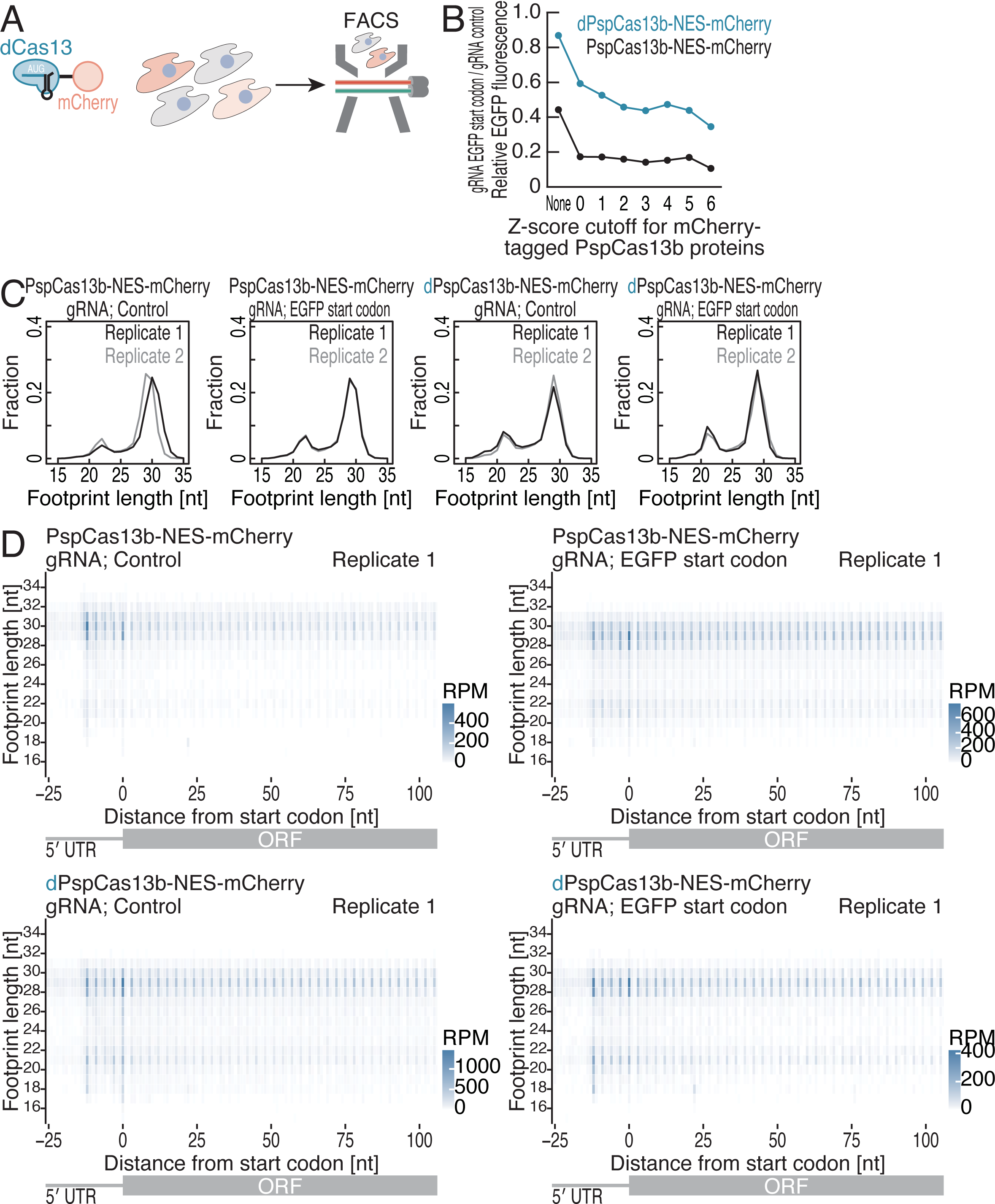
Characterization of ribosome profiling data with flow cytometry-sorted cells, related to Figure 3. (A) Schematic of an experimental procedure for measuring EGFP (reporter) and mCherry (PspCas13b variant) expression. PspCas13b variant expression was ensured by cell sorting based on mCherry expression. (B) Relative EGFP fluorescence plotted against the PspCas13b variant expression level. Signals from the fused mCherry were used to assess PspCas13b expression. Data from 1 × 10^4^ cells were used for the analysis, with subpopulation means determined with the indicated Z score cutoffs. (C) Footprint length distributions under the indicated conditions. (D) Metagene plots for the 5′ ends of ribosome footprints for the indicated samples. The data were aligned to the start codons. The color scales for the reads are shown. RPM, reads per million mapped reads.

**Figure S4.**
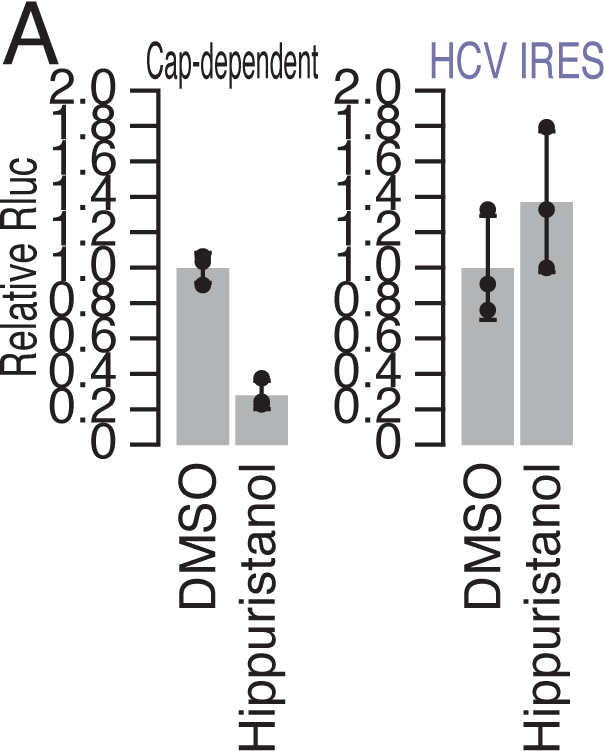
Characterization of HCV IRES-driven translation, related to Figure 4. (A) Relative Rluc luminescence from the indicated reporter mRNAs after 1 μM hippuristanol treatment. The mean (gray bar), s.d. (black line), and individual replicates (n = 3, black point) are shown.

**Figure S5.**
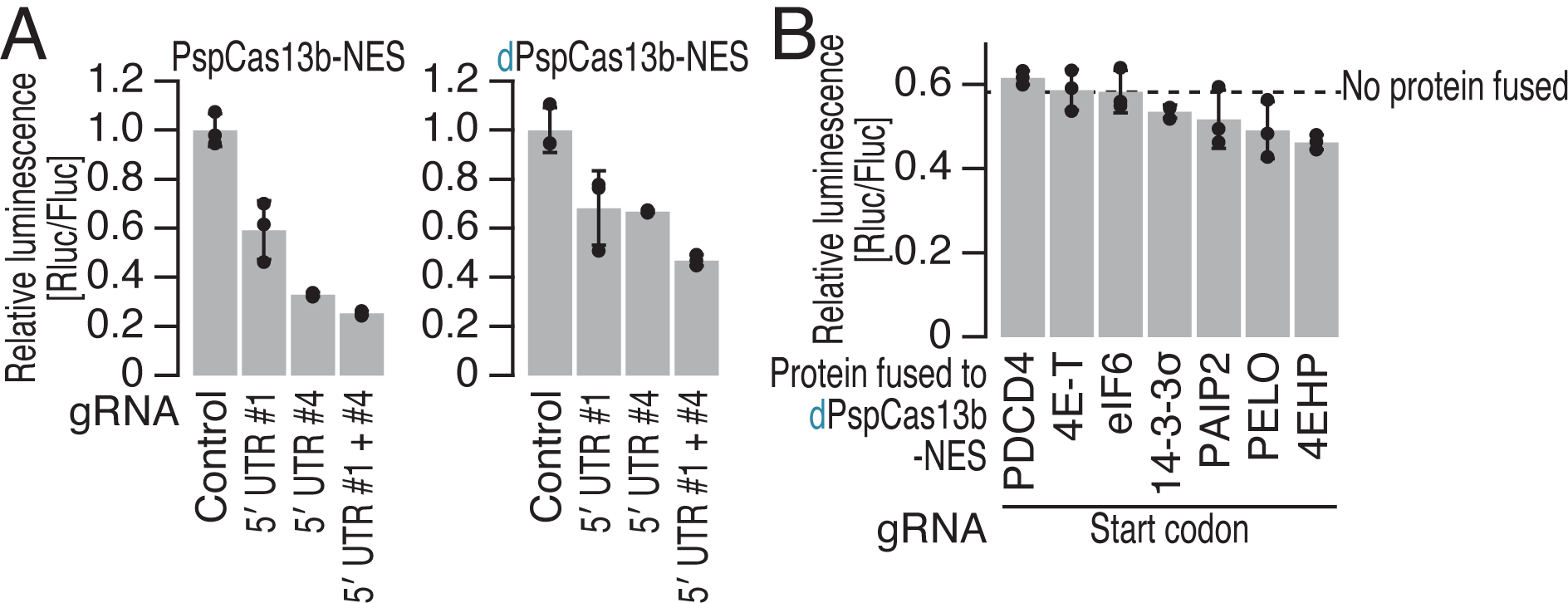
Options for improving CRISPRδ efficacy, related to Figure 5. (A) Relative Rluc luminescence with respect to Fluc luminescence was calculated to quantify the knockdown activity of PspCas13b-NES and dPspCas13b-NES using the gRNAs shown in Figure 1G. (B) Relative Rluc luminescence with respect to Fluc luminescence was calculated to quantify the knockdown activity of dPspCas13b-NES alone and fused to translational repressors using a gRNA targeting the start codon of the Rluc reporter mRNA. In A and B, the mean (gray bar), s.d. (black line), and individual replicates (n = 3, black point) are shown.

**Table S1. gRNA expression vectors prepared in this study**. Each plasmid is listed with target plasmid or transcript, gRNA name, spacer sequence, DR sequence, and primer sets used for its construction.

## Notes

### Competing Interest Statement

The authors have declared no competing interest.

